# Mitochondrial transfer between breast cancer cells promotes ROS-dependent proliferation

**DOI:** 10.64898/2026.08.26.747325

**Authors:** Noah M. Bressler, Taylor Z. Stevens, Minna Roh-Johnson

## Abstract

Previously, we showed that macrophages transfer mitochondria to breast cancer cells, promoting proliferation in acceptor cancer cells. Transferred mitochondria were depolarized and accumulated reactive oxygen species (ROS), and the mitochondrial transfer-induced proliferation was dependent upon ROS signaling (Kidwell et al. 2023). Our unexpected findings supported a model in which transferred mitochondria act as a signal for proliferation in acceptor cancer cells rather than a direct source of increased bioenergetics. It remains unclear whether this unexpected signaling mechanism is unique to macrophages as the donor cell, or whether this mechanism applies to mitochondrial transfer between other cells within the tumor microenvironment. Here, we show that highly metastatic cancer cells transfer mitochondria to weakly metastatic cancer cells. These transferred mitochondria are depolarized, accumulate ROS, and promote ROS-dependent proliferation in acceptor cancer cells. Furthermore, we specifically attribute this proliferative phenotype to the transfer of mitochondria, as when we isolate mitochondria from highly metastatic cells and apply these purified mitochondria directly to weakly metastatic cells in culture, acceptor cancer cells that internalize the purified mitochondria exhibit increased proliferation in a ROS-dependent manner. These findings support mitochondrial transfer within the breast tumor microenvironment as a signaling axis for proliferation, regardless of donor cell identity.

## INTRODUCTION

The intercellular transfer of mitochondria serves a vital role in stress responses (Spees et al. 2006; Hayakawa et al. 2016; Mistry et al. 2019; Li et al. 2024), tissue homeostasis (Nicolas-Avila et al. 2020; Levoux et al. 2021; Xu et al. 2026), and modulating cellular function (Baldwin et al. 2024; Court et al. 2024). In cancer, mitochondrial transfer promotes proliferation (Kheirandish-Rostami et al. 2020; Roushandeh et al. 2020; Kidwell et al. 2023; Watson et al. 2023; Kuo et al. 2024; Ikeda et al. 2025), metastasis (Ippolito et al. 2019; Goliwas et al. 2023; Frisbie et al. 2024; Hoover et al. 2025), and drug resistance (Moschoi et al. 2016; Sansone et al. 2017; Nakhle et al. 2023; Watson et al. 2023; Del Vecchio et al. 2024) in acceptor cancer cells. While the observed changes in acceptor cell behavior were initially posited to occur through a direct enhancement of energy supplied by transferred mitochondria (Spees et al. 2006; Islam et al. 2012; Jackson et al. 2016; Moschoi et al. 2016; Marlein et al. 2017; Saha et al. 2022), more recent studies, including from our group, have challenged this notion by reporting the transfer of dysfunctional or depolarized mitochondria (Nicolas-Avila et al. 2020; Crewe et al. 2021; Lampinen et al. 2022; Kidwell et al. 2023). Given that, mitochondrial membrane polarization is essential for numerous mitochondrial functions, including oxidative phosphorylation, the transfer of depolarized mitochondria suggests that these mitochondria likely have impaired ATP production. However, the transfer of depolarized mitochondria has been observed under both physiological (Rosina et al. 2022; Xia et al. 2026) and pathological conditions, including cancer (Kidwell et al. 2023; Terasaki et al. 2026). Furthermore, in many mitochondrial transfer studies, only a small amount of mitochondria is transferred to acceptor cells, raising the question of how the transfer of a small amount of mitochondria could meaningfully enhance energy production in the acceptor cell, particularly when the acceptor cell already has a functioning endogenous mitochondrial network (Liao et al. 2024; Xia et al. 2026). Together, these studies challenge the dogma that proposes transferred mitochondria are functional and simply restore and enhance energy for the cell.

We previously showed that macrophages transfer mitochondria to breast cancer cells in vitro and in vivo (Kidwell et al. 2023). These transferred mitochondria were not degraded and did not fuse with the endogenous host mitochondrial network of acceptor cancer cells, suggesting transferred mitochondria constitute a unique and distinct subpopulation in acceptor cells. Surprisingly, transferred macrophage mitochondria were depolarized and accumulated reactive oxygen species (ROS). We also found that ROS generation was sufficient to activate the ERK pathway, promoting proliferation of acceptor cancer cells. This work suggested that ROS accumulation at transferred mitochondria and downstream signal propagation led to increased proliferation of acceptor cancer cells, providing a potential explanation as to how a small amount of transferred mitochondria could have a larger functional impact in acceptor cells. However, this form of mitochondrial transfer-induced cell-cell communication was unexpected at the time and led to the question of whether this signaling mechanism is specific to macrophage mitochondrial transfer or more generally applicable to mitochondrial transfer in breast cancer.

To address this gap in understanding, we changed the donor cell to another breast cancer cell and asked whether mitochondria transferred from highly metastatic breast cancer cells to weakly metastatic breast cancer cells are depolarized and regulate acceptor cancer cell proliferation in a ROS-dependent manner. Breast cancer patients show a high degree of intratumoral heterogeneity, with cancer cells exhibiting variability in mutational status and therefore, metastatic potential (Li et al. 2021; Luond, Tiede, and Christofori 2021; Angus et al. 2019). Numerous studies have shown that malignant cells confer increased metastatic behavior and proliferation to less aggressive cancer cells through the transfer of cytoplasmic contents (Zomer et al. 2015), mitochondrial DNA (Ishikawa et al. 2008; Tan et al. 2015; Dong et al. 2017; Rabas et al. 2021), and intact mitochondria (Jing et al. 2024), but the mechanisms by which transferred cytoplasmic contents, and specifically transferred mitochondria, exert their effects on acceptor cell behavior remain poorly understood.

By studying both spontaneously-transferred and artificially-transferred (bath-applied) mitochondria from highly metastatic to weakly metastatic breast cancer cells, we found that transferred mitochondria are depolarized and accumulate ROS in weakly metastatic acceptor cells. Furthermore, weakly metastatic acceptor cells that receive mitochondria from highly metastatic cancer cells exhibit increased proliferation. We also found, similar to our previous studies with macrophages, this mitochondrial transfer-induced proliferation is dependent on ROS. Together, these data support a model in which tumor cells transfer depolarized mitochondria to breast cancer cells, promoting proliferation in acceptor cells through ROS-dependent signals.

## RESULTS

### Highly metastatic breast cancer cells transfer mitochondria to weakly metastatic breast cancer cells

We previously reported that macrophages transfer depolarized mitochondria to breast cancer cells, promoting the proliferation of acceptor cancer cells in a ROS-dependent manner (Kidwell et al. 2023). We hypothesized that this signaling mechanism is not unique to mitochondrial transfer from macrophages but is a more generalizable signaling mechanism within the breast cancer tumor microenvironment. To test this hypothesis, we used the highly metastatic human breast cancer line, MDA-MB-231 (231 cells), as the donor cells and the weakly metastatic human breast cancer line, MCF7, as the acceptor cells. To track the transfer of mitochondria from 231 cells to MCF7 cells, we stably expressed a mitochondrially-localized mEmerald (mito-mEm) fluorescent protein in the 231 donor cells, and we stably expressed a membrane-localized mCherry (mCh-CAAX) in the MCF7 acceptor cells (**Figure 1a**). We co-cultured these cell lines for 24 hours at a donor-to-acceptor cell ratio of 3-to-1. After 24 hours, we observed mEm+ mitochondria in mCh+ MCF7 acceptor cells. Using 3D analysis, we confirmed that these mitochondria are inside of MCF7 acceptor cells (**Figure 1b**). To quantify mitochondrial transfer from 231 cells to MCF7 cells, we used flow cytometry to determine the percentage of mCh+ MCF7 cells with mEm+ mitochondria (**Figure 1c**). To reliably discriminate between donor and acceptor cells, we stably co-expressed a nuclear localized blue fluorescent protein (NLS-BFP) in the 231 donor cells. The NLS-BFP allowed us to exclude NLS-BFP+ cells from our analyses and only quantify mitochondrial transfer from 231 cells to MCF7 cells rather than transfer events in the opposite direction or cell-cell fusion events (**Supplementary Figure 1**). We tested three different donor-to-acceptor (D:A) ratios at 1:1, 3:1, or 5:1 and found across all conditions, the proportion of mitochondrial transfer acceptor cells was consistently ∼1% with flow cytometric analysis of 24 hour co-cultures (**Figure 1d**), compared to 0.2% in control samples in which 231 mito-mEm cells were added to mCherry-CAAX MCF7 cells at time zero (t = 0 hr control). For comparison, an average of 0.84% ± 0.10 231 cells receive mitochondria from macrophages in 24hr co-cultures (Kidwell et al. 2023). While these mitochondrial transfer rates were consistent, we hypothesized that based on the small amount of mitochondria visualized in acceptor cells by live imaging (**Figure 1b**), we were likely excluding a large percentage of acceptor cells that contain transferred mEm+ mitochondria when analyzing mitochondrial transfer by flow cytometry. Thus, we also quantified mitochondrial transfer by live imaging. Using the 3:1 D:A ratio, we co-cultured 231 mito-mEm cells with MCF7 mito-RFP cells, randomly acquired images across the imaging dishes, and then quantified the percentage of MCF7 mito-RFP cells that contained mEm+ mitochondria (**Figure 1e**, left). We used z-stack analysis and orthogonal projections to confirm that the mEm+ mitochondria were inside the MCF7 mito-RFP cells. With this analysis, we found that the rate of mitochondrial transfer was 17.67% ± 4.33% (**Figure 1e**, right). Furthermore, in all cases, we do not observe co-localization of transferred mEm+ mitochondria with RFP+ mitochondria, suggesting transferred mitochondria do not fuse with the host mitochondrial network of MCF7 acceptor cells. Together, these results show that highly metastatic breast cancer cells transfer mitochondria to weakly metastatic cells and comprise a distinct subpopulation of mitochondria in the acceptor cell.

**Figure 1.**
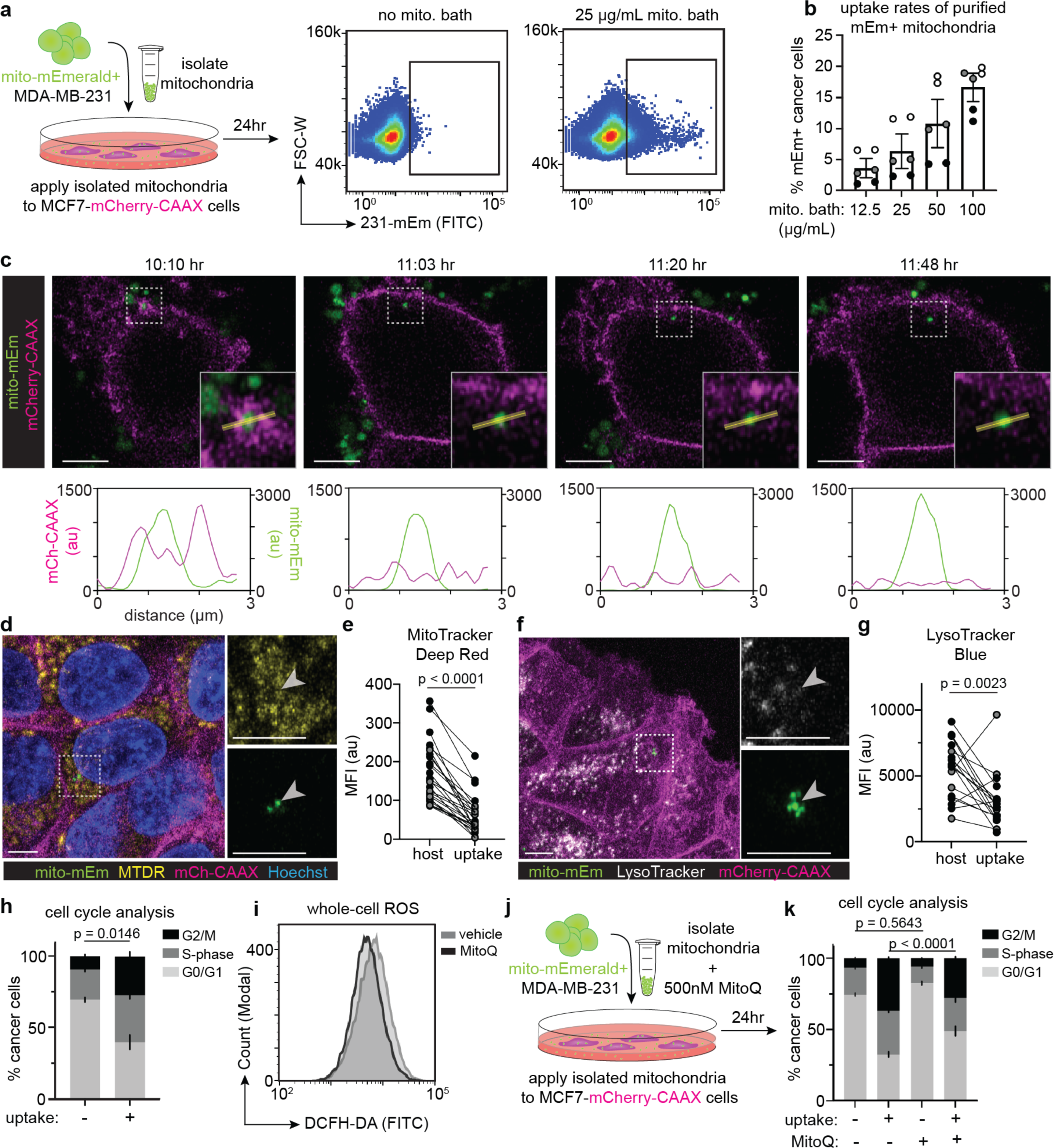
Highly metastatic MDA-MB-231 cancer cells transfer mitochondria to weakly metastatic MCF7 cancer cells. **(a)** Left: Co-culture schematic of highly metastatic MDA-MB-231 cells expressing a mitochondrially-localized mEmerald tag (mito-mEm) co-cultured with weakly metastatic MCF7 cells expressing a membrane-localized mCherry tag (mCh-CAAX). Right: A max-projection representative confocal microscopy image of a MDA-MB-231 mito-mEm (green) and MCF7 mCh-CAAX (magenta) 24 hour co-culture. **(b)** A single z-slice representative image of a mitochondrial transfer event. An MCF7 cell (magenta) with transferred mitochondria (green, arrowhead) from an MDA-MB-231 cell. Using 3D orthogonal view projection analysis (1μm z-step size) and the mCh-CAAX membrane tag (magenta), the transferred mitochondria are confirmed to be inside the MCF7 acceptor cell (right). **(c)** Representative flow cytometry plots with an “acceptors” gate around the MCF7 acceptor cells in 24hr co-cultures of MDA-MB-231 NLS-BFP + mito-mEm cells and MCF7 mito-RFP cells. A time-zero control (t = 0 hr; left) was used to define an “acceptors” gate on 0.2% of MCF7 cells. This gate was then applied unchanged to the 24 hour co-cultures to quantify mitochondrial transfer. **(d)** Quantification of mitochondrial transfer rates in a 24-hour co-culture when plated at a 1:1, 3:1, and 5:1 donor-to-acceptor cell ratio (D:A). Each data point represents a technical replicate; each color represents a biological replicate (N = 3 experiments, 2 technical replicates per experiment). **(e)** 3:1 co-cultures of MDA-MB-231 mito-mEm cells and MCF7 mito-RFP cells were imaged after 24 hours to quantify the percentage of transfer acceptors (left) in each field of view. Quantification of these fields of view are plotted (right) with each data point representing a field of view and each color representing biological replicates (511 cells across 72 fields of view; N = 3 experiments). For all images, scale bars = 10µm.

### Mitochondria transferred from highly metastatic donor cells are depolarized and promote proliferation of weakly metastatic acceptor cells

In our previous work, we showed that macrophage mitochondrial transfer to breast cancer cells promotes breast cancer proliferation (Kidwell et al. 2023). Thus, to determine if mitochondrial transfer from highly to weakly metastatic breast cancer cells also promotes acceptor cell proliferation, we used flow cytometric approaches. By evaluating DNA content and Ki67 expression, a marker of proliferation (**Figure 2a; Supplemental Figure 2a**), we classified cells into phases of the cell cycle. We found significant increases in the percent of acceptor cells in the G2 and Mitotic (M) phases of the cell cycle, compared to their sister cells in the same co-culture that did not accept mitochondria (**Figure 2a,b**), suggesting enhanced proliferation. To determine whether these cells were actively dividing, we analyzed only the M-phase and found a significantly higher percentage of acceptor cells in the M-phase of the cell cycle compared to sister cells that did not accept mitochondria (**Supplemental Figure 3a**). To further support these findings, we also performed an orthogonal proliferation assay and evaluated EdU labeling to quantify the percentage of cells actively undergoing DNA replication (**Figure 2c**). With this approach, we again found that a significantly higher percentage of MCF7 cells that accepted mitochondria from 231 cells underwent DNA replication compared to MCF7 cells that did not accept mitochondria in the same co-culture (**Figure 2d**). Together, these results suggest that weakly metastatic MCF7 cells that receive mitochondria from highly metastatic 231 cells exhibit increased proliferative capacity.

**Figure 2.**
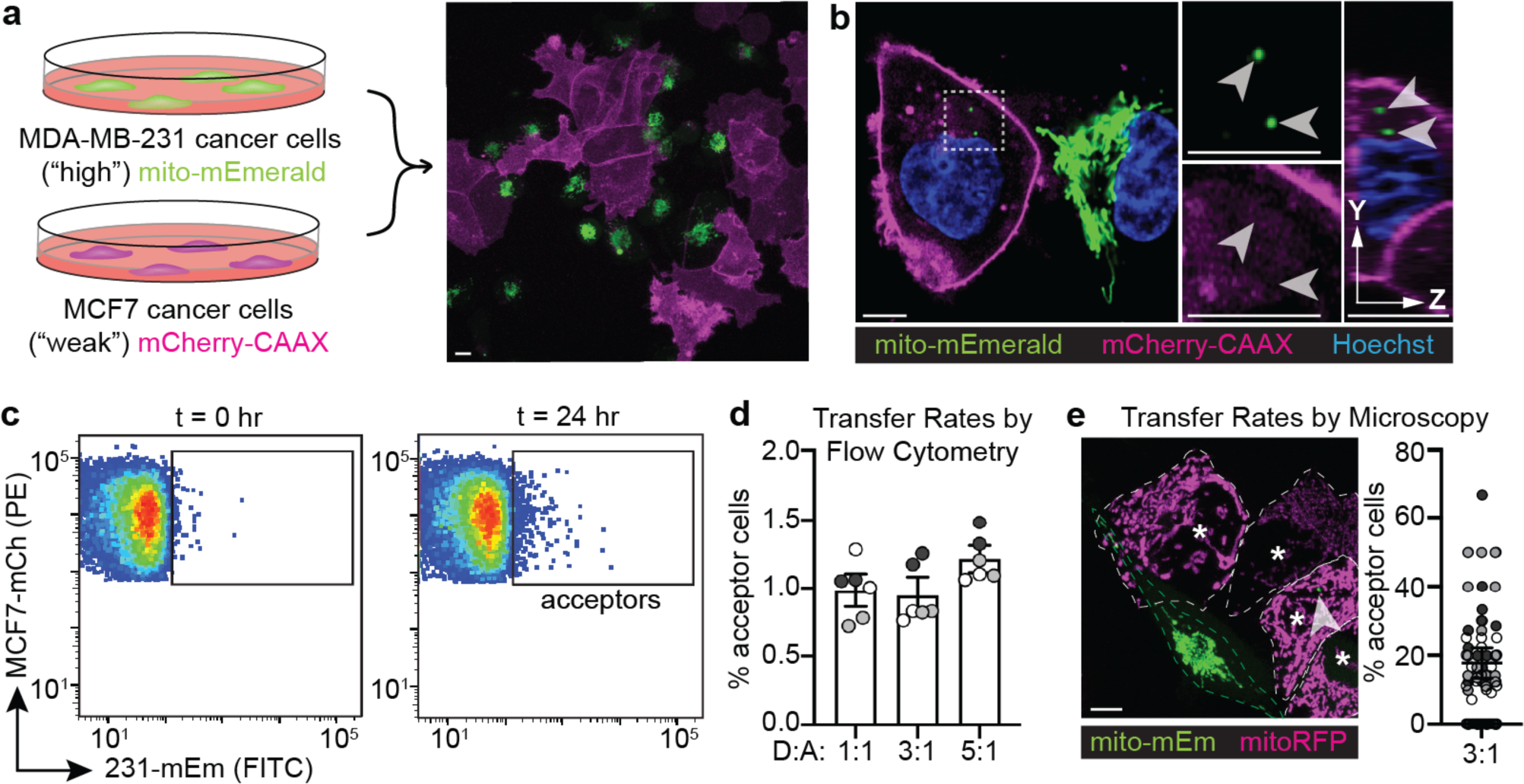
Mitochondria transferred from highly metastatic cancer cells are depolarized and promote proliferation in weakly metastatic acceptor cells. **(a)** Representative flow cytometry plots of MCF7 non-acceptor cells (left) and MCF7 acceptor cells (right) within the same co-culture. Using DAPI for total DNA content (x-axis) and Ki67 as a marker of proliferation (y-axis), cells were gated in G0/G1, S-phase, or G2/M-phase. **(b)** Quantification of cell cycle states of MCF7 non-acceptor cells vs. MCF7 acceptor cells. Statistics are only shown for G2/M-phase comparisons depicted in black (N = 3 experiments, 2 technical replicates per experiment). **(c)** Representative flow cytometry histograms of EdU fluorescence in MCF7 non-acceptor cells (left) and MCF7 acceptor cells (right) within the same co-culture. EdU-and EdU+ gates were determined using an unlabeled control co-culture, where EdU was added, but the fluorescent conjugate was omitted. **(d)** Quantification of the percentage of EdU- and EdU+ cells in MCF7 non-acceptor cells and MCF7 acceptor cells (N = 3 experiments, 3 technical replicates per experiment). **(e)** Representative image of an MCF7 acceptor cell (magenta) labeled with MitoTracker Deep Red (MTDR; yellow). Zoomed images (right) show transferred mitochondria (green; arrowhead) lacking MTDR signal. Asterisks denote individual MCF7 cells; arrowheads denote mitochondrial transfer events. **(f)** Quantification of the mean fluorescence intensity (MFI) of MTDR signal at transferred mitochondria (transfer) relative to the mean fluorescence intensity (MFI) of MTDR signal of the endogenous mitochondrial network (host) within each cell. Each pair of data points represents an individual MCF7 acceptor cell with each color representing a biological replicate (23 cells across N = 3 experiments). **(g)** Representative image of an MCF7 acceptor cell (magenta) labeled with LysoTracker Blue (white). Zoomed images (right) show transferred mitochondria (green, arrowhead) lacking LysoTracker Blue signal. **(h)** Quantification of the MFI of LysoTracker Blue signal at transferred mitochondria (transfer) relative to the MFI of LysoTracker Blue signal inside the acceptor cell (host). Each pair of data points represents an individual MCF7 acceptor cell with each color representing a biological replicate (17 cells across N = 3 experiments). For all panels with error bars, standard error of the mean (SEM) is shown. For all images, scale bars = 10µm. Paired t-test (**b,d**), Wilcoxon signed-rank test (**f,h**). au, arbitrary units.

To next determine whether acceptor MCF7 cells exhibited a dose-dependent increase in proliferation based on the amount of transferred mEm+ mitochondria the cell received, we divided the acceptor MCF7 cell population into “high” mitochondrial acceptors (received more transferred mEm+ mitochondria; acceptor cells within the top 50% of expression) versus “low” mitochondrial acceptors (received less transferred mEm+ mitochondria; acceptor cells within the bottom 50% of expression) (**Supplemental Figure 3b**). We observed no statistically significant difference in the percentage of acceptor cells in the G2/M phase of the cell cycle between these two populations (**Supplementary Figure 3c),** suggesting that there was no dose-dependent relationship between the amount of transferred mitochondria and the proliferation rate that we could detect.

To determine the functional status of the transferred mitochondria, we stained co-cultures with MitoTracker Deep Red (MTDR), a dye that specifically accumulates in mitochondria that have membrane potential (**Figure 2e**). We quantified the fluorescence intensity of the MTDR signal at transferred mitochondria (transfer) and compared this value to the average MTDR signal of the endogenous mitochondrial network of the acceptor cell (host). We found that the MTDR signal at transferred mitochondria was dramatically reduced compared to the average signal of the host cell mitochondrial network, indicating that transferred mitochondria are depolarized in MCF7 acceptor cells (**Figure 2f**).

We then asked whether the transferred mitochondria are depolarized because they are encapsulated in acidic vesicles and are being subjected degradation. We stained co-cultures with LysoTracker Blue, a dye that specifically accumulates in acidic vesicles, including lysosomes (**Figure 2g**). We quantified the LysoTracker Blue signal at transferred mitochondria and found that mitochondria transferred to MCF7 cells did not co-localize with LysoTracker Blue (**Figure 2h**), indicating that these mitochondria were not located inside acidic vesicles. Together with the MTDR results, these results suggest that transferred mitochondria are depolarized but are not being subjected to degradation in acidic vesicles.

The 231 and MCF7 cell lines arise from two different breast cancer patients and are genetically distinct. Thus, we sought to determine whether mitochondrial transfer occurred from highly metastatic cancer cells to weakly metastatic cancer cells with the same genetic backgrounds. We used “matched pairs” in which the metastatic line was derived from the initial primary tumor. E0771 murine mammary adenocarcinoma cells spontaneously arose in a C57BL/6J mouse and exhibit relatively low metastatic potential (Casey, Laster, and Ross 1951). The highly metastatic counterpart line, E0771.lmb, was generated from a spontaneous metastatic lung nodule isolated from an orthotopically injected mouse (Johnstone et al. 2015). Thus, we generated E0771.lmb cells (highly metastatic) expressing mito-mEm and co-cultured them with E0771 (weakly metastatic) expressing mito-RFP (**Figure 3a**). After 24hr, we found that 1.13 ± 0.16% of E0771 cells contained transferred mEm+ mitochondria, compared to 0.2% in t = 0 hr controls by flow cytometry (**Figure 3b,c**). Similar to what we observed with the human breast cancer cells, mEm+ mitochondria transferred from E0771.lmb highly metastatic cells to E0771 weakly metastatic cells are depolarized (**Figure 3d,e**). We also found that E0771 weakly metastatic cells that received mitochondria from E0771.lmb highly metastatic cells exhibit enhanced proliferative capacity (**Figure 3f,g; Supplemental Figure 3d**). Taken together, these results suggest that highly metastatic breast cancer cells transfer mitochondria to weakly metastatic cancer cells. The transferred mitochondria remain distinct from the host mitochondrial network, are depolarized, and acceptor cells that receive transferred mitochondria exhibit increased proliferation.

**Figure 3.**
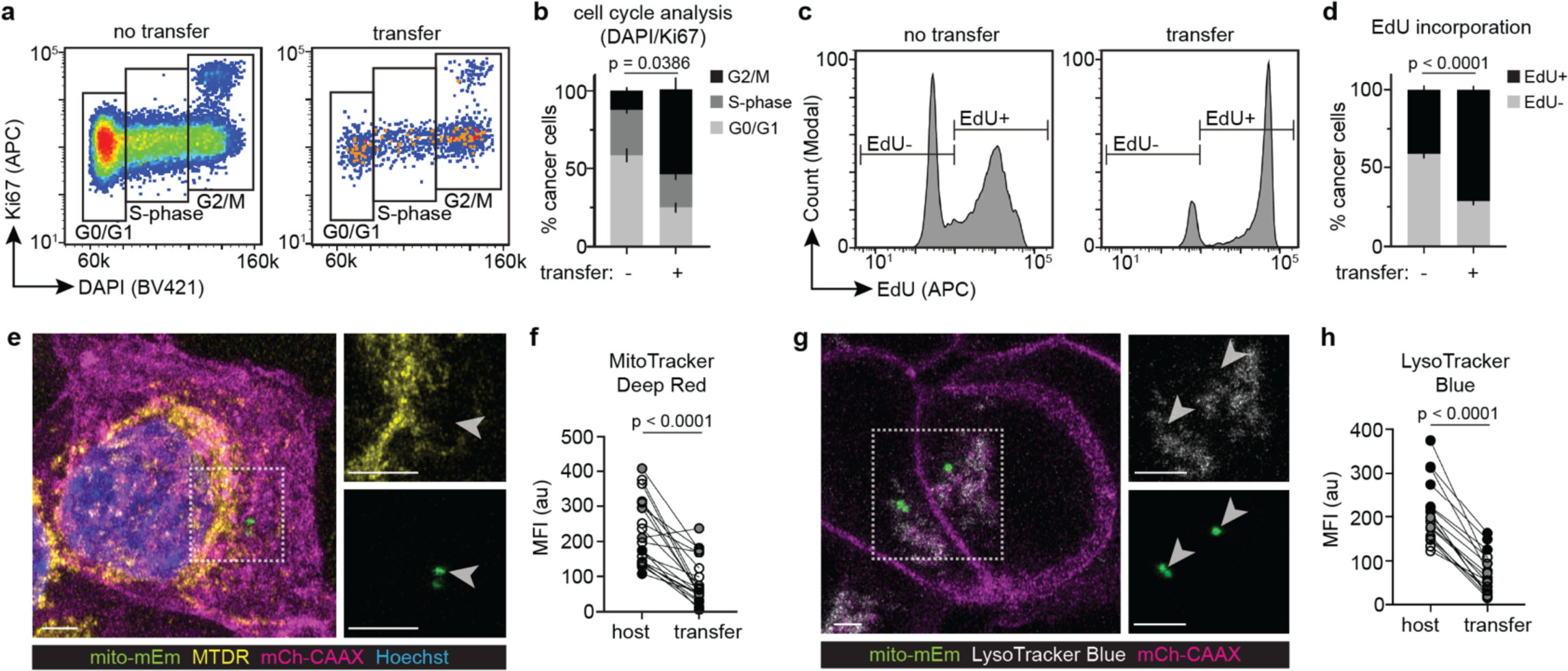
Weakly metastatic murine adenocarcinoma cells that accept mitochondria from highly metastatic murine adenocarcinoma cells exhibit increased proliferation. **(a)** Left: Co-culture schematic of highly metastatic E0771.lmb cells expressing a mitochondrially-localized mEmerald tag (mito-mEm) co-cultured with weakly metastatic E0771 cells expressing a mitochondrially-localized RFP tag (mito-RFP). Right: A max-projection representative confocal microscopy image demonstrates that the E0771.lmb cells (green) and E0771 cells (magenta) are interspersed after 24 hours of co-culturing (right) with examples of mitochondrial transfer (arrowheads). **(b)** Representative flow cytometry plots with a gate around the E0771 acceptor cells. A time-zero (t = 0 hr) control (left) is used to set a gate on 0.2% of the E0771 population. The 24-hour co-cultures (t = 24 hr, right) use the same gate as the t = 0 hr control to quantify the percentage of E0771 acceptor cells. The right boundary of the gate did not include the brightest RFP+/mEm+ double-positive cells to avoid counting cell fusion events as mitochondrial transfer events. **(c)** Quantification of the percentage of E0771 cells that received mitochondrial transfer in a 3:1 co-culture after 24 hours. Each data point represents a technical replicate and each color depicting a biological replicate (N = 3 experiments with 2 technical replicates per experiment). **(d)** Representative image of an E0771 acceptor cell (magenta) labeled with MitoTracker Deep Red (MTDR; yellow). Zoomed images (right) show transferred mitochondria (green; arrowheads) lacking MTDR signal. **(e)** Quantification of the mean fluorescence intensity (MFI) of MTDR signal at transferred mitochondria (transfer) relative to the MFI of MTDR signal inside the acceptor cell (host). Each pair of data points represents an individual E0771 acceptor cell with each color representing a biological replicate (15 cells across N = 1 experiment). **(f)** Representative flow cytometry plots of E0771 non-acceptor cells (left) and E0771 acceptor cells (right) within the same co-culture. Using DAPI for total DNA content (x-axis) and Ki67 as a marker of proliferation (y-axis), cells were gated in G0/G1, S-phase, or G2/M-phase. **(g)** Quantification of cell cycle states of E0771 non-acceptor cells vs. E0771 acceptor cells. Statistics are only shown for G2/M-phase comparisons depicted in black (N = 3 experiments with 2 technical replicates per experiment). Error bars represent SEM, and for all images, scale bars = 10µm. Wilcoxon signed-rank test (**e**), paired t-test (**g**).

### Mitochondria from highly metastatic cells accumulate ROS and promote ROS-dependent proliferation in weakly metastatic acceptor cells

Given that transferred mitochondria from highly metastatic cells to weakly metastatic cells are depolarized (**Figure 2e,f**), we next asked whether mitochondria transferred from highly metastatic cells to weakly metastatic cells accumulate ROS, as we observed with transferred macrophage mitochondria (Kidwell et al. 2023). We expressed a redox-sensitive biosensor targeted to the mitochondria, mito-Grx1-roGFP2, in MCF7 weakly metastatic cells as a readout of the mitochondrial glutathione redox state. In its native, reduced form, this biosensor exhibits maximal fluorescence when excited with 488nm light. In the presence of ROS, glutathione is oxidized, and the biosensor reaches maximal excitation with 405nm light (**Figure 4a**). Thus, we used the redox state of glutathione as a proxy for ROS levels. For these studies, we generated 231 donor cells expressing RFP-tagged mitochondria. We then co-cultured 231 mito-RFP cells with MCF7 cells expressing mito-Grx1-roGFP2. We found that after 24 hours, there was significant enrichment of the oxidized form of glutathione at transferred mitochondria compared to the host mitochondrial network (**Figure 4b**). We quantified the oxidized:reduced forms of glutathione at transferred mitochondria (transfer) compared to the host mitochondrial network of each acceptor cell (host) and found significant accumulation of oxidized glutathione at transferred mitochondria (**Figure 4c**). These results suggest that transferred mitochondria accumulate ROS in acceptor cancer cells.

**Figure 4.**
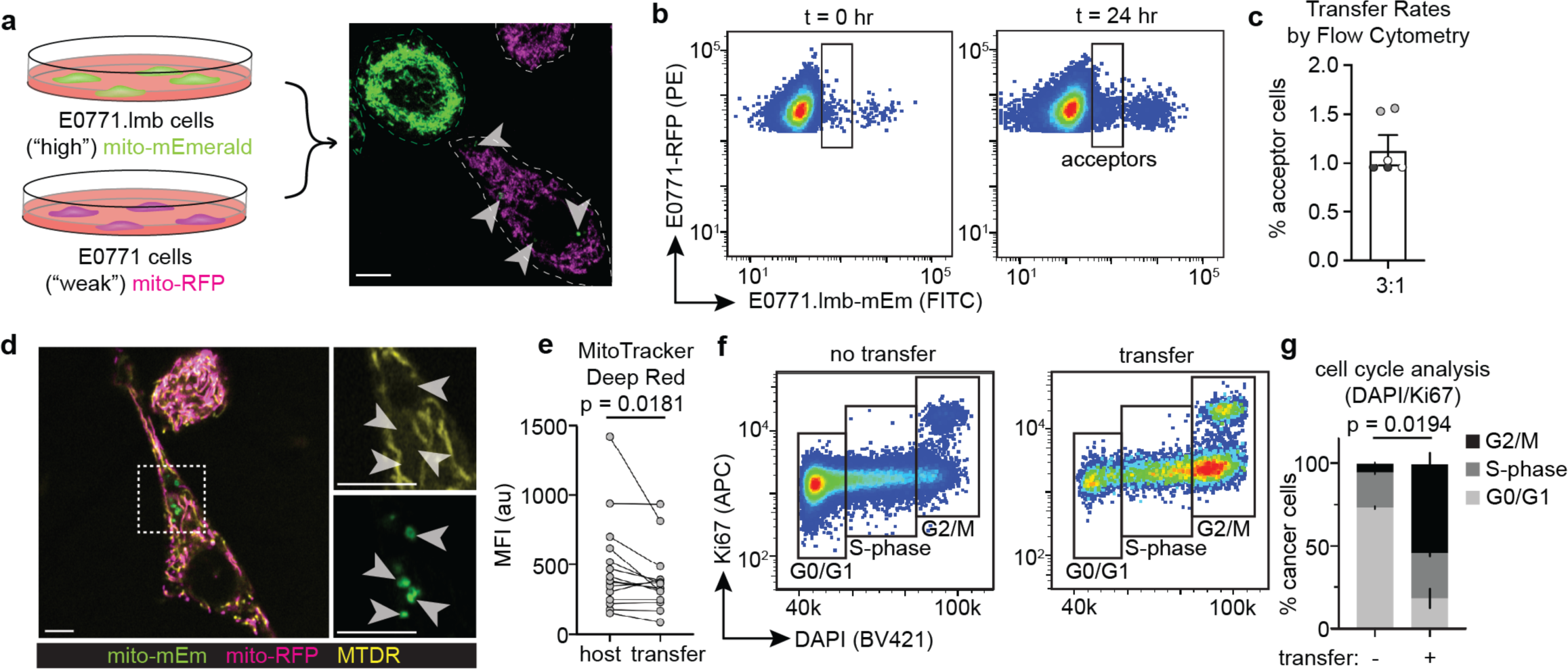
Mitochondria transferred from highly metastatic cells accumulate reactive oxygen species in weakly metastatic acceptor cells. **(a)** Schematic of the mitochondrially-localized redox-sensitive biosensor (mito-Grx1-roGFP2) expressed in MCF7 cells. Grx1 couples glutathione oxidation to the oxidation state of roGFP2. In the native, reduced form, the biosensor exhibits peak excitation at 488nm. In the oxidized form, the biosensor exhibits peak excitation at 405nm. **(b)** Representative image of an MCF7 acceptor cell (white outline, green) with transferred RFP+ mitochondria (magenta, arrowheads). Zoomed in panels (right) show each individual channel: the biosensor in its reduced state (green; top left), the biosensor in its oxidized state (yellow; top right), transferred mitochondria (magenta; bottom left), and a ratiometric image of the oxidized-to-reduced fluorescence intensity ratio (purple-to-yellow gradient; bottom right). **(c)** Quantification of the oxidized-to-reduced ratio of transferred mitochondria (transfer) relative to the average oxidized-to-reduced ratio of the endogenous mitochondrial network inside the acceptor cell (host). Each pair of data points represents a single MCF7 acceptor cell with each color representing a biological replicate (17 cells across N = 4 experiments). For all images, scale bar = 10µm. Wilcoxon signed-rank test (**c**).

### Mitochondria purified from highly metastatic cells are internalized by weakly metastatic cells and promote proliferation

To determine if the enhanced proliferation phenotype observed in MCF7 acceptor cells is mediated by the receipt of mitochondria versus other factors, we purified mEm+ mitochondria from 231 highly metastatic cells and bath applied them to MCF7 weakly metastatic cells (**Figure 5a**). At a concentration of 12.5μg/mL, we found that 3.60 ± 1.56% of MCF7 weakly metastatic cells take up purified mEm+ mitochondria after 24 hours. Furthermore, MCF7 cells exhibited a dose-dependent increase in uptake to 16.63 ± 2.26% when 100μg/mL of purified mEm+ mitochondria were added (**Figure 5b**). Using timelapse imaging, we found that purified mEm+ mitochondria initially decorated the outside of MCF7 cells. MCF7 cells then took up small amounts of mitochondria, similar to the levels that we observe in the co-culture systems (**Figure 5c**). Exogenous mitochondria taken up by MCF7 cells are first enclosed in a membrane, and then the membrane is no longer detected ∼1-2 hours later (**Figure 5c**). We also determined that purified mitochondria taken up by MCF7 cells are depolarized (**Figure 5d,e**) and do not colocalize with acidic vesicles (**Figure 5f,g**). We subsequently asked if purified mitochondria promote proliferation in accepter cancer cells, similar to the spontaneous transfer observed in our co-cultures. We found that MCF7 cells that have taken up exogenous mitochondria have a significantly greater percentage of cells in the G2/M and M-only phases of the cell cycle compared to MCF7 cells that did not take up exogenous mitochondria in the same culture (**Figure 5h; Supplemental Figure 3e**). These results suggest that the increased proliferative capacity exhibited by acceptor MCF7 cells is due to the receipt of exogenous mitochondria.

**Figure 5.**
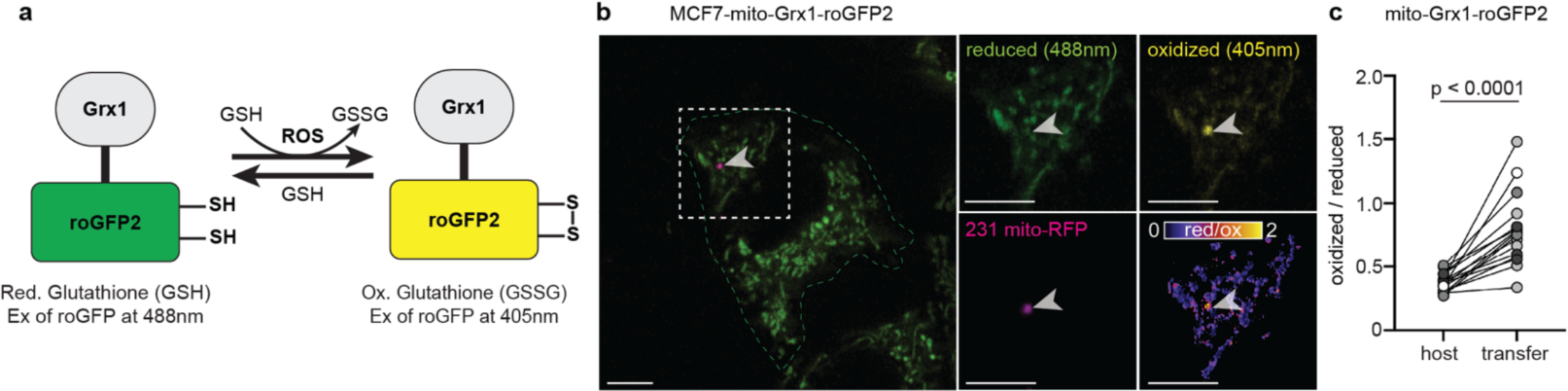
Mitochondria purified from highly metastatic cells are depolarized and promote ROS-dependent proliferation in weakly metastatic acceptor cells. **(a)** Left: Schematic of purification and application of MDA-MB-231 mEm+ mitochondria to MCF7 mCh-CAAX cells in culture. Right: Representative flow cytometry plots (right) of MCF7 cells that did not receive purified mitochondria (no mito. bath; left) or MCF7 cells that received 25µg/mL of purified mEm+ mitochondria (25µg/mL mito. bath; right). **(b)** Quantification of the percentage of MCF7 cells that internalized purified mitochondria in a dose-dependent manner. Each data point represents a technical replicate, and each color represents a biological replicate (N = 3 experiments, 2 technical replicates per experiment). **(c)** Top: Images from a timelapse recording of an MCF7 cell (magenta) internalizing purified mitochondria (green). Each image is a single z-plane. Times listed above each image depicts the time after addition of the purified mitochondria. Zoomed insets are shown in the bottom right corner. Bottom: Line scan analysis of mEm and mCh signal corresponding to each image above. A line of 5 pixels wide and 2.7µm long was used (yellow line in the insets above). Left y-axes depict pixel intensity of mEm (mitochondria) signal. Right y-axes depict mCh (membrane) signal. **(d)** Representative image of an MCF7 acceptor cell (magenta) labeled with MitoTracker Deep Red (MTDR; yellow). Zoomed images (right) show internalized mitochondria (green; arrowhead) lacking MTDR signal. **(e)** Quantification of the mean fluorescence intensity (MFI) of MTDR signal at internalized mitochondria (uptake) relative to the MFI of MTDR signal of the endogenous mitochondrial network of the acceptor cell (host). Each pair of data points represents an individual MCF7 acceptor cell with each color representing a biological replicate (25 cells across N = 2 experiments). **(f)** Representative image of an MCF7 acceptor cell (magenta) labeled with LysoTracker Blue (white). Zoomed images (right) show internalized mitochondria (green, arrowhead) lacking LysoTracker Blue signal. **(g)** Quantification of the MFI of LysoTracker Blue signal at internalized mitochondria (uptake) relative to the MFI of LysoTracker Blue signal inside the acceptor cell (host). Each pair of data points represents an individual MCF7 acceptor cell with each color representing a biological replicate (18 cells across N = 2 experiments). **(h)** Cell cycle analysis of MCF7 non-acceptor cells (left; -) and MCF7 acceptor cells (right; +) that have not and have internalized purified mitochondria, respectively. Statistics are only shown for G2/M-phase comparisons depicted in black (N = 5 experiments, 2 technical replicates per experiment). **(i)** Flow cytometric histograms of DCFH-DA fluorescence in MCF7 cells after 4 hr treatment with 500nM MitoQ (dark gray) or vehicle (light gray). **(j)** Schematic of purification and application of MDA-MB-231 mEm+ mitochondria with 500nM MitoQ or vehicle to MCF7 mCh-CAAX cells in culture. **(k)** Cell cycle analysis of MCF7 cells that internalized purified mEm+ mitochondria (uptake +) compared to sister cells that did not (uptake -) with treatment of vehicle (MitoQ -) or 500nM MitoQ (MitoQ +), a mitochondrial-targeted antioxidant. 50µg/mL of purified mEm+ mitochondria were applied. Statistics are only shown for G2/M-phase comparisons depicted in black (N = 4 experiments, 2 technical replicates per experiment). Error bars represent SEM, and all scale bars = 10µm. Wilcoxon signed-rank test (**e,g**), paired t-test (**h**), two-way ANOVA (Fisher’s LSD post-hoc; **k**).

### Mitochondrial transfer-induced cell proliferation is ROS-dependent

Our data suggest that highly metastatic cancer cells transfer depolarized mitochondria to weakly metastatic cancer cells. Depolarized mitochondria accumulate reactive oxygen species, and weakly metastatic acceptor cancer cells that accept mitochondria from highly metastatic cancer cells exhibit increased proliferative capacity. To directly test whether ROS accumulation is necessary for the increased proliferative capacity exhibited by MCF7 acceptor cells, we quenched mitochondrial ROS with a mitochondrially-targeted antioxidant, MitoQ (mitoquinone mesylate). We confirmed a reduction in ROS upon MitoQ treatment by staining MCF7 cells with DCFH-DA, a fluorescent probe that measures intracellular ROS (**Figure 5i**). We then asked whether treating MCF7 cells with MitoQ was sufficient to attenuate the proliferation phenotype of MCF7 cells that take up exogenous mitochondria. To achieve a high number of “uptake-positive” cells for downstream analysis, we added 50μg/mL of purified 231 mEm+ mitochondria to MCF7 cells and then treated cells with MitoQ or vehicle control (**Figure 5j**). In acceptor MCF7 cells, we observed a significant reduction in the percentage of cells in the G2/M and M-only phases of the cell cycle when treated with MitoQ compared to vehicle (**Figure 5k**, compare black bars 2 & 4; **Supplemental Figure 3f**). Importantly, this ROS-dependent function was specific to acceptor MCF7 cells, as non-acceptor MCF7 cells (ie. MCF7 cells that did not take up exogenous mEm+ mitochondria) did not exhibit a change in the percentage of cells in the G2/M and M-only phases of the cell cycle upon MitoQ treatment (**Figure 5k**; compare black bars 1&3; **Supplemental Figure 3f**). Together, these data indicate that transferred mitochondria accumulate ROS and promote ROS-dependent proliferation in weakly metastatic acceptor cells, similar to breast cancer cells that receive transferred mitochondria from macrophages.

## DISCUSSION

While recent studies have provided new insight into the cellular processes and dynamics regulating mitochondrial transfer in cancer (Zhang et al. 2023; Cangkrama et al. 2025; Novak et al. 2025), the effects of mitochondrial transfer on acceptor cell biology and behavior remain an ongoing debate. Initial studies of mitochondrial transfer posited that acceptor cells directly receive increased energy through the transfer of functional mitochondria (Spees et al. 2006; Cho et al. 2012; Islam et al. 2012; Ahmad et al. 2014; Jackson et al. 2016; Ippolito et al. 2019; Saha et al. 2022). However, our previous publication suggests that the mitochondria transferred to cancer cells are depolarized and induce acceptor cell behavior through signaling mechanisms (Kidwell et al. 2023). This surprising finding raised the question of whether the observed signaling mechanism was unique to mitochondrial transfer from macrophages to cancer cells, or whether transferred mitochondria serve as a signal for acceptor cell proliferation between other donor-acceptor pairs within the tumor microenvironment. In this work, we find that mitochondria transferred from highly metastatic to weakly metastatic cells are depolarized, accumulate ROS, and promote proliferation through ROS-dependent signaling. Thus, we propose that mitochondrial transfer serves as a signal to induce acceptor cell proliferation in various donor-acceptor pairs within the tumor microenvironment, including mitochondrial transfer from macrophages to cancer cells and between cancer cells.

Several questions arise from this work. First, why are transferred mitochondria depolarized in the acceptor cells? Related, is there a specific subpopulation of mitochondria that are preferentially transferred by the donor cell or preferentially taken up by the acceptor cell? Recent publications have identified distinct subpopulations of mitochondria within cells that differ in their structure, function, and proteomic profile (Benador et al. 2018; Schulte et al. 2023; Ryu et al. 2024; Glover et al. 2026). The highly dynamic, specialized, and heterogenous nature of mitochondria and their subpopulations raise the possibility that only certain mitochondria are selectively transferred. Furthermore, phenotypic changes in acceptor cells may be dependent upon the specific mitochondria that are transferred.

Another question that arises from this work is what is the fate of transferred mitochondria in the acceptor cell? Under homeostatic conditions, dysfunctional mitochondria in a cell are typically subjected to lysosomal degradation, in a process known as mitophagy, or fusion with healthy mitochondria (Ashrafi and Schwarz 2013; Quintana-Cabrera and Scorrano 2023). A recent publication demonstrated that mitochondria transferred from cancer cells are enriched in a de-ubiquitinating enzyme, USP30, allowing the transferred mitochondria to escape degradation in acceptor cells (Ikeda et al. 2025). This finding provides a potential explanation of why the transferred mitochondria are not degraded in acceptor cells. Studies using genetically encoded fluorescent proteins have also suggested fusion between transferred mitochondria and acceptor cell host mitochondria (Cowan et al. 2017; Huang et al. 2024; Ikeda et al. 2025). However, we have not observed fusion in our imaging-based experiments, including this work and our previous publication (Kidwell et al. 2023). It is possible that the fate of transferred mitochondria is determined by the donor-acceptor cell pair. In a third, alternative fate of dysfunctional mitochondria, multiple groups have suggested that post-mitotic cells can shuttle mitochondria to surrounding cells for subsequent degradation, in a process coined as “trans-mitophagy” (Davis et al. 2014; Morales et al. 2020; Nicolas-Avila et al. 2020; Lampinen et al. 2022; Xia et al. 2026). Because we rarely observe transferred mitochondria co-localize with lysosomal markers in our studies, it is unlikely that such a process is occurring when mitochondria are transferred between cancer cells. Yet, a recent publication suggests that a subpopulation of transferred mitochondria accumulate in previously uncharacterized “mitochondrial degradation bodies” (MDBs) (Glover et al. 2026), warranting further investigation into MDBs as a possible explanation for how transferred mitochondria escape canonical lysosomal degradation and remain distinct from the acceptor cell’s host mitochondrial network. Additional research is needed to elucidate the regulation and specificity of mitochondrial transfer in regard to both donor and acceptor cell biology.

## MATERIALS AND METHODS

### Cell culture of cell lines

The human cell line MDA-MB-231 (HTB-26; RRID:CVCL_0062) and the murine cell line E0771 (CRL-3461; RRID:CVCL_GR23) were directly purchased from American Type Culture Collection and cultured according to their recommendations. The human cell line MCF7 (HTB-22; RRID:CVCL_0031) and the highly metastatic murine cell line variant E0771.lmb (CRL-3405; RRID:CVCL_B0A2) were gifts from Keren Hilgendorf and cultured according to American Type Culture Collection recommendations. Cell lines were authenticated through STR profiling, and all cultured cell lines were subjected to mycoplasma testing every 6 months using the Universal Mycoplasma Detection Kit (30-1012K, ATCC). Base media used was DMEM, high glucose (11965118, ThermoFisher) and 10% heat-inactivated fetal bovine serum (FBS; A5256501, ThermoFisher). All cell lines were kept in culture for no more than 25 passages total.

### Generation of mito-mEmerald, mito-RFP, mCherry-CAAX and NLS-BFP stable cell lines

The mito-mEmerald plasmid (Addgene #174542; RRID:Addgene_174542) and mito-RFP plasmid (Addgene #174543; RRID:Addgene_174543) were generated as previously described (Kidwell et al. 2023). In brief, the cytochrome oxidase subunit VIII mitochondrial targeting sequence was fused to either *mEmerald* or *TagRFP-T and* introduced into a pLKO.1 plasmid backbone with an accessible multiple cloning site (pLKO.1_MCS) to generate lentivirus. The pLenti-mCherry-CAAX (mCherry-CAAX) plasmid was purchased from Addgene (#129285; RRID:Addgene_129285) and used to generate lentivirus. The lenti-NLS-mTagBFP2 (NLS-BFP) was purchased from Addgene (#216128; RRID:Addgene_216128) and used to generate lentivirus. Then, stable lines were generated through lentiviral transduction. For transduction, approximately 50,000 cells were plated into one well of a 6-well plate directly with the appropriate lentivirus supernatant diluted 1:5 in DMEM complete media with a final concentration of 10μg/mL polybrene (Sigma, TR-1003-G). After 48-72hr, cells were expanded, and multiclonal populations were flow sorted for appropriate levels of fluorescent protein expression. All other transgenic cell lines were generated as outlined in subsequent sections.

### Lentivirus production

pLKO.1_MCS and pLenti plasmids containing the appropriate transgene were used to generate lentivirus as outlined previously (Johnson et al. 2020). In brief, HEK 293 FT cells (RRID:CVCL_6911) in 15cm plates were transfected with PEI-max (24765, Polysciences) and plasmids for pCMV-VSV-G (Addgene #8454; RRID:Addgene_8454), psPax2 (Addgene #12260; RRID:Addgene_12260), and transgene cassettes. The following day, cells were washed and grown for an additional 36hr in fresh media. Supernatants were harvested, passed through 0.45μm filters, and used fresh. Lentiviral supernatants were used to transduce cell lines as outlined in “generation of mito-FP” section unless otherwise noted.

### Flow cytometry

The following flow cytometry machines were used: a BD FACS Aria (equipped with 4 Lasers: 405, 488, 561, 640; RRID:SCR_018934) referred to as the Aria, or a BD LSR Fortessa (5 Lasers: UV, 405, 488, 561, 640; RRID:SCR_018655) referred to as the Fortessa. Technical details per experiment type are listed below.

### Stable line generation

Cells were enzymatically dissociated using trypsin-EDTA (25200056, ThermoFisher) and resuspended in buffer consisting of 0.5% Bovine Serum Albumin (BSA; Sigma, A9418) in DPBS (14190250, ThermoFisher). Cells were sorted according to fluorescence intensity on the Aria and collected in the appropriate media containing 0.5% Pen/Strep (15140122, ThermoFisher).

### Mitochondrial transfer quantification via flow cytometry

Cells were enzymatically dissociated using trypsin-EDTA, washed with ice-cold DPBS, and resuspended in ice-cold DPBS for analysis on the Fortessa. The background level of mEmerald fluorescence was set at 0.2% as described previously (Kidwell et al. 2023). In brief, this background level was based on a co-culture control where donor cells were not transduced with mito-mEmerald. This gate was defined by FACS-isolating co-cultures of mCherry-CAAX acceptor cells and mito-mEm donor cells and determining a gate that accurately isolated acceptor cells containing mEm+ mitochondria. A 0.2% threshold predominantly isolated acceptor cells containing mEm+ mitochondria, as confirmed by microscopy (Kidwell et al. 2023). To exclude cell fusions from mitochondrial transfer quantification, we stably expressed a nuclear localized blue fluorescent protein (NLS-BFP) in 231 donor cells, and we excluded cells that were positive for all markers (BFP+/mCh+/mEm+ triple-positive population; **Figure 1c,d; Supplemental Figure 1**). In co-cultures where DAPI/Ki67 cell cycle analysis was conducted, the NLS-BFP tag was not expressed due to the use of DAPI. Instead, the “transfer-positive” gate only extended to ∼10^3.5^ to exclude the brightest mEm+ cells from cell cycle analysis, due to the high likelihood that these were fusion events as we have previously shown (**Figure 2a,b**; **Figure 3f,g; Supplemental Figure 2**). The right-most edge of this “transfer-positive” gate was verified against mEm+ donor cell monocultures to ensure that no more than 1% of the monocultured mEm+ donor cells would be included in the gate. Additionally, for flow cytometry-based experiments, mCherry-CAAX expression was eventually swapped for mito-RFP expression in acceptor cell lines due to the high propensity of generating mCh+/mEm+ double-positive cells in 24hr co-cultures; expressing mito-RFP in the acceptor cell lines generated less double-positive cells. For all experiments using the matched-pair murine cell lines, E0771.lmb mito-mEm and E0771 mito-RFP were used (**Figure 3a-c**).

### Mitochondrial transfer quantification via microscopy

MDA-MB-231 expressing mito-mEmerald and MCF7 cells expressing mito-RFP were combined at a 3:1 ratio and plated at an approximate density of 400,000 cells directly onto 35mm glass bottom dishes (FD35-100, World Precision Instruments). After 24 hours, co-cultures were imaged at random locations throughout the 35mm dishes. Images were then analyzed for the presence of MCF7 acceptor cells with mEm+ mitochondria. In each image captured, the percentage of mEm+ MCF7 acceptor cells were quantified. Using the 3D “orthogonal view” function in FIJI (RRID:SCR_002285; **Figure 1b**), mEm+ mitochondria were determined to be inside or outside MCF7 cells.

### Quantification of DNA content and Ki67

Co-cultures were enzymatically dissociated with trypsin-EDTA then fixed and permeabilized on ice for 15 min using the eBioscience Foxp3/Transcription Factor Staining Buffer Set (00-5523-00, ThermoFisher) according to the manufacturer’s instructions. Cells were stained with an APC-conjugated Ki67 antibody (APC-Ki67; 17-5699-42, ThermoFisher; RRID:AB_2573218) at 1:40 for 20 min followed by a 3 μM DAPI (D9542, Sigma) solution for 10 min. Cells were resuspended in cold DPBS for analysis on the Fortessa. The background level of mEm fluorescence in the “transfer-positive” gate was set at 0.2% (see “mitochondrial transfer quantification” section). The highest mEm+ cell population was not included in the final “transfer-positive” gate and subsequent analysis to avoid including fusion events, or mEm+ and RFP+ cells that are stuck together, which are commonly seen via confocal microscopy of 24 hr co-cultures. Thus, the upper limit of the “transfer-positive” gate was determined using either MDA-MB-231 mito-mEm or E0771.lmb mito-mEm monocultures.

### Quantification of EdU labeling

EdU labeling assays were performed according to the manufacturer’s instructions included in the Click-iT™ EdU Alexa Fluor™ 647 Flow Cytometry Assay Kit (C10419, ThermoFisher). In brief, co-cultures were treated with 10µM EdU 12hr prior to collection for flow cytometry analysis. Upon collection for flow cytometry analysis, cells were pelleted and resuspended in Click-iT™ fixative and incubated in the dark for 15min. Next, cells were permeabilized in the dark for 15min. Then, cells were treated with 1x Click-iT™ reaction cocktail and incubated in the dark for 30min. After washing, pelleting, and resuspending cells in ice-cold DPBS, cells were analyzed on the Fortessa. Co-cultures with EdU added but no reaction cocktail were used to determine EdU positivity in experimental co-cultures.

### Live imaging of co-cultures with cell-permeable dyes

Imaging was performed using a Zeiss LSM 880 with AiryScan technology (Carl Zeiss, Germany; RRID:SCR_020925) and a 63x/1.4 NA oil objective. Images were acquired using the AiryScan Fast mode. For all live imaging, cells were maintained at 37°C, 5% CO2 with an on-stage incubator. For human co-cultures, MDA-MB-231 cells and MCF7 cells stably expressing appropriate transgenes were mixed at a 3:1 ratio and plated at an approximate density of 400,000 cells directly onto 35mm glass bottom dishes (FD35-100, World Precision Instruments) for all live imaging experiments. Mouse co-cultures were plated in the same format and density. Duration of co-cultures are indicated in main text or figure legend.

For detection of nuclear DNA, Hoechst 33342 (B2261, Sigma) was diluted into culture media to a final concentration of 5μg/mL. After 10min at 37°C, cells were washed with warm DPBS, and warmed complete media was replaced before imaging.

For detection of mitochondrial membrane potential with MitoTracker Deep Red (MTDR; M22426, ThermoFisher) at a final concentration of 25nM and incubated at 37°C for 30min. Following incubation, cells were washed with warm DPBS, and warmed complete media was replaced before imaging. To quantify MTDR signal at transferred mitochondria, the Otsu thresholding algorithm was used to define the area of mEm+ mitochondria in acceptor cells and create a region of interest (ROI) where the mean gray value of MTDR signal was quantified. This value was then paired to the average MTDR signal inside the cell; the same Otsu thresholding algorithm was used to define the MTDR+ ROI inside the acceptor cell, and the mean gray value of the entire MTDR+ ROI was quantified.

For detection of lysosomes and acidic vesicles, LysoTracker Blue (L7525, ThermoFisher) was diluted to a final concentration of 100nM in serum-free DMEM media and incubated at 37°C for 30min. Following incubation, cells were washed with warm DPBS, and warmed complete media was replaced before imaging. To quantify LysoTracker Blue signal at transferred mitochondria, the Otsu thresholding algorithm was used to define the area of mEm+ mitochondria in acceptor cells and create an ROI where the mean gray value of LysoTracker Blue signal was quantified. This value was then paired to the average LysoTracker Blue signal inside the cell; the same Otsu thresholding algorithm was used to define the LysoTracker Blue+ ROI inside the acceptor cell, and the mean gray value of the entire LysoTracker Blue+ ROI was quantified.

For detection of ROS, 2’,7’-dichlorofluorescin (DCFDA; ab113851, Abcam) was diluted to 20μM into warmed DMEM without phenol-red. After a 30min incubation at 37 °C, cells were washed with ice-cold DPBS and resuspended in ice-cold DPBS for flow cytometry analysis. As a positive control for DCFH-DA sensitivity to changes in ROS, a 4hr treatment of 100μM *tert*-Butyl hydroperoxide (TBHP) was used and compared to untreated samples. Then, the mean fluorescence intensity was compared between untreated and MitoQ-treated samples to verify antioxidant activity after a 4hr treatment.

### ROS biosensor line generation, imaging, and quantification

MCF7 cells were transfected with the following plasmids: pLPCX mito-Grx1-roGFP2 (Gutscher et al. 2008) (Addgene, plasmid #64977; RRID:64977) using the Lipofectamine™ 3000 Transfection Kit (L3000008, ThermoFisher) according to the manufacturer’s instructions. Cells were allowed 3-7 days to recover and then sorted on the Aria for expression. Biosensor expressing MCF7 cells were co-cultured with mito-RFP expressing 231 cells for 24hr and imaged on the Zeiss LSM 880. Cells were sequentially imaged (per z-plane) for the presence of transferred RFP+ mitochondria (Ex. 561nm, Em. BP 570-620nm + LP 645nm) and the biosensor in its reduced (Ex. 488nm, Em. BP 420-480nm + BP 495-550nm) and oxidized (Ex. 405nm, Em. BP 420-480nm + 495-550nm) form. Images were initially processed using Zen software (see “image analysis” section) and further analysis was performed using FIJI as indicated previously (Morgan, Sobotta, and Dick 2011). pLPCX mito-Grx1-roGFP2 was a gift from Tobias Dick.

### Mitochondrial isolation and bath application

150-200 x 10^6^ mito-mEm expressing 231 cells were pelleted by centrifugation for 5 min at 300 *g*. Pellets were resuspended in 1 mL of modified CP-1 buffer (Chappell and Perry 1954) (100mM KCl, 50mM Tris-HCl, 2mM EGTA, 1x protease inhibitor, pH 7.4) and incubated on ice for 30min to induce cell swelling. Suspended cells were Dounce homogenized 80-100 times in a Potter-Elvehjem PTFE pestle and glass tube (Sigma, P7734). The glass tube was washed with 1 mL of isotonic buffer (225mM mannitol, 75mM sucrose, 1mM EGTA, 10mM HEPES, 1x protease inhibitor, pH 7.4) and transferred to a 2mL microcentrifuge tube. Cell homogenates were centrifuged at 700 *g* for 10 min at 4°C to pellet and remove nuclei. The supernatant was then centrifuged at 3,000*g* for 15min at 4°C, and the crude mitochondrial pellet was washed with 1 mL of ice-cold isotonic buffer. This step was repeated two additional times at 12,000*g* to remove unwanted cellular material. The resulting mitochondrial pellet was resuspended in 100-500uL of ice-cold isotonic buffer, and relative mitochondrial protein concentrations were determined via a standard BCA protein concentration assay (ThermoFisher, 23225). 50μg/mL of mitochondria were applied to pre-plated MCF7 cells for 24hr unless otherwise noted. After mitochondrial incubation, cells were thoroughly washed with DPBS to remove any un-internalized mitochondria. The percentage of cell that internalized mEm+ mitochondria was quantified on the Fortessa, and cell cycle analyses were conducted as described in “quantification of DNA content and Ki67”.

For live time-lapse imaging of MCF7 cells treated with isolated mitochondria: 20-30μg/mL of isolated mitochondria were bath applied for 4hr to remove excessive amounts of un-internalized mitochondria. Cells were imaged for 12-16hr on the Zeiss LSM 880 AiryScan confocal microscopy as described in “live cell imaging”.

### Line scan analysis of internalized mitochondria

To determine the presence and persistence of acceptor cell membrane around uptaken purified mitochondria, we performed line scan analysis on timelapse images captured while an MCF7-mCh-CAAX cell internalized an exogenous mEm+ mitochondrion (**Figure 5c**). Using a 5 pixel-wide line drawn across the uptaken mitochondrion and the “plot profile” function in FIJI, mCh and mEm fluorescence were tracked across various representative images of the timelapse, revealing the persistence of mEm signal and gradual decline of mCh signal flanking the mEm signal.

### Drug treatment: Mitoquinol mesylate (MitoQ)

To quench mitochondrial ROS, mitoquinol mesylate (MitoQ; 89950, Cayman Chemical) was formulated at 100mM in 100% DMSO, diluted into warm complete media to a concentration of 100μM, and further diluted to a working concentration of 500nM. MCF7 cells were treated for the duration of 24hr as described in “Mitochondrial isolation and bath application”, and cells were harvested for proliferative capacity analyses (see quantification of Ki67 and DNA content section). MitoQ aliquots were stored at -20°C, remained protect from light, and never underwent a freeze-thaw cycle. As a positive control for ROS induction and DCFH-DA sensitivity, cells were treated with 100μM *tert*-Butyl hydroperoxide (TBHP) diluted directly into warm DMEM without phenol red for 4hr prior to collection for flow cytometry analysis. To verify a reduction in whole-cell ROS levels upon MitoQ treatment, cells were treated for 4hr with 500nM MitoQ. In the final 30 min of treatment, DCFH-DA was added to a final concentration of 20μM. To determine if MitoQ treatment was sufficient to reduce proliferation in MCF7 acceptor cells, 500 nM MitoQ was added simultaneously with purified mitochondria. Cell cycle analysis via flow cytometry was performed 24hr after addition of the drug and mitochondria.

### Data and materials availability

All data are available in the main text or in the Supplementary Data.

### Image analysis

All images taken with the AiryScan detector on the Zeiss LSM 880 were subjected to deconvolution using the Zen Black software (Carl Zeiss; RRID:SCR_018163) with “auto” settings (referred to as AiryScan processed). Maximum intensity projections of selected z-planes were generated using FIJI software (Schindelin et al. 2012). Linear adjustments to the brightness and contrast were made using FIJI. Images were cropped and panels were assembled using Adobe Photoshop and Illustrator, respectively (Adobe, Inc; RRID:SCR_014199; RRID:SCR_010279).

### Graphical representations and statistical analysis

All graphs were generated using Prism software (v10, GraphPad; RRID:SCR_002798). All graphs show mean with standard error of the mean. Statistical analyses were performed using Prism. Statistical tests used and p-values are indicated in each figure legend. Flow cytometry data and representations were analyzed and generated using FlowJo software (v10.10, BD; RRID:SCR_008520). When comparing cell populations originating from the same well (**Figure 2b,d**; **Figure 3g**; **Figure 5h**), paired statistics were used. When comparing cell populations originating from different wells (**Figure 5k**), un-paired statistics were used. Two-tailed paired t-tests were used when we compared mean values of paired data with normal distribution, and two-tailed Wilcoxon signed-rank tests were used when the paired data were not normally distributed. Two-way ANOVA was utilized when comparing how two independent variables influence a dependent variable followed by Fisher’s LSD multiple comparison post-hoc test. Sex as a biological variable was not assessed in this study due to the use of human cell lines. All statistical methodologies were performed under the guidance of biostatistician, Dr. Kenneth M. Boucher.

## ACKNOWLEDGEMENTS

We thank all members of the Roh-Johnson lab for helpful discussions and edits to this manuscript; Dr. Keren Hilgendorf and Abigail Jackson for providing cell lines; and Dr. Kenneth M. Boucher for help with statistics. We also thank the Huntsman Cancer Institute Cancer Center Shared Resources; the University of Utah Flow Cytometry Core (RRID:SCR_012210) for technical assistance; and the University of Utah Cell Imaging Core (RRID:SCR_023496) for the use of confocal microscopes. This work was supported by the National Cancer Institute R37CA247994 and the Department of Defense Breast Cancer Research Program BC191176 (to MRJ); National Cancer Institute R37CA247994-04S1 and the Intermountain Post-baccalaureate Research Education Program (IM-PREP) R25GM144253 (to TZS); and the Office of The Director of the National Institutes of Health S10OD026959 and National Cancer Institute 5P30CA042014-24 (to the Flow Cytometry Core).

## AUTHOR CONTRIBUTIONS

Conceptualization, NMB, TZS, MRJ; Methodology, NMB, TZS, MRJ; Investigation, NMB, TZS, MRJ; Writing, NMB, MRJ; Writing – Reviewing & Editing, NMB, TZS, MRJ; Funding Acquisition, TZS, MRJ; Supervision, MRJ.

