## Supplemental Figures for "Mitochondrial transfer between breast cancer cells promotes ROS-dependent proliferation"

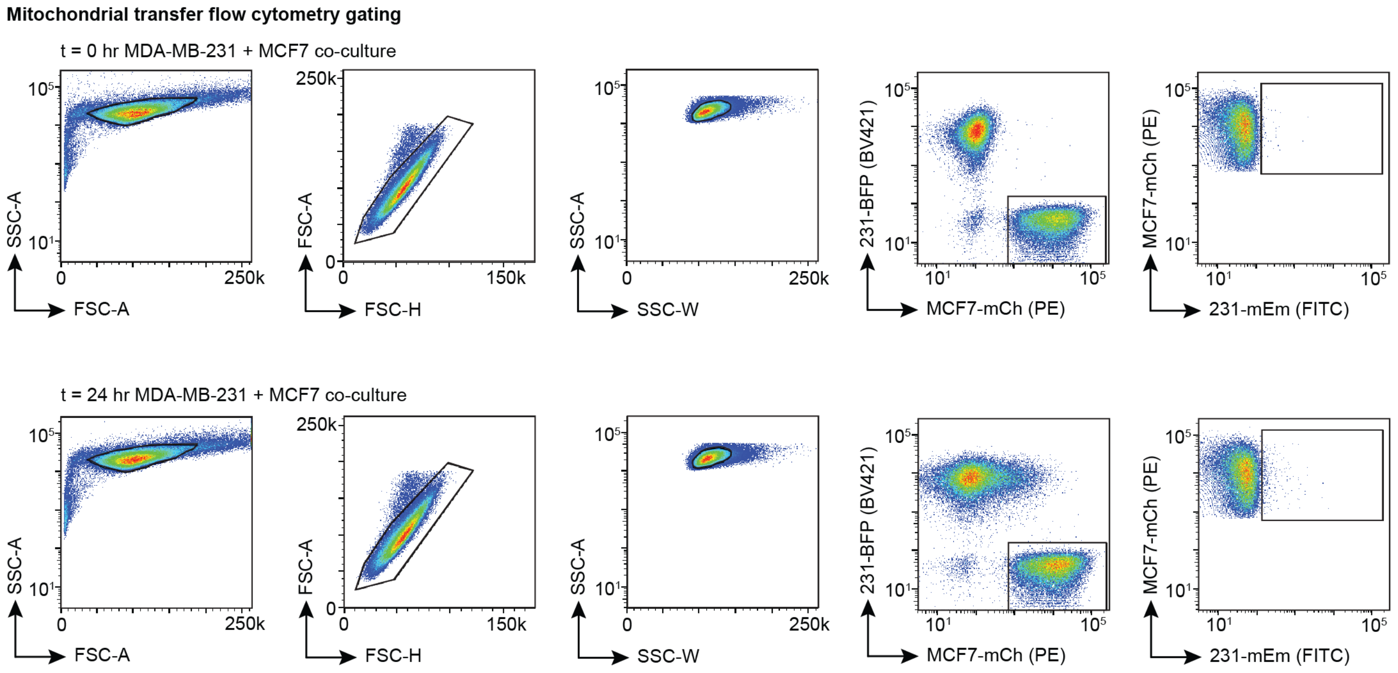


**Supplemental Figure 1. Flow cytometry gating scheme to isolate MCF7 mCh-CAAX single cells in co-cultures with MDA-MB-231 mito-mEm + NLS-BFP cells.**

After gating for single cells, mCh+/BFP- cells were classified as MCF7 mCh-CAAX cells. To quantify the percentage of mEm+ MCF7 acceptor cells in t = 24 hr co-cultures, the final "acceptor" gate was set first to 0.2% in the t = 0 hr control co-culture (top). The final "acceptor" gate was then applied to the t = 24 hr co-cultures for quantification (bottom).

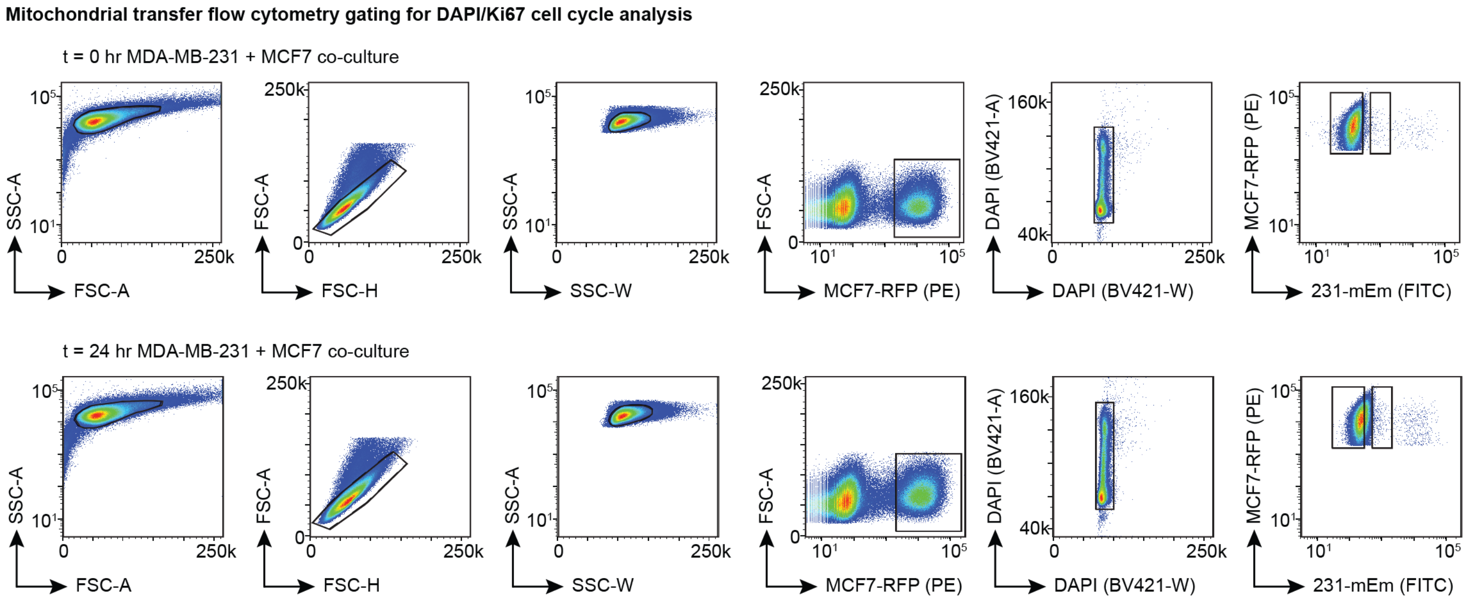


**Supplemental Figure 2. Flow cytometry gating scheme to isolate MCF7 mito-RFP cells in co-culture with MDA-MB-231 mito-mEm cells.**

To discriminate mEm+ acceptor MCF7 cells from mEm- non-acceptor MCF7 cells in t = 24 hr co-cultures, the final “acceptor” gate was set first to 0.2% in the t = 0 hr control co-culture (top). The final “acceptor” gate was then applied to the t = 24 hr co-cultures (bottom) for analysis of cell cycle states using DAPI and Ki67. A “non-acceptor” gate was also used to analyze cell cycle states of non-acceptor MCF7 cells.

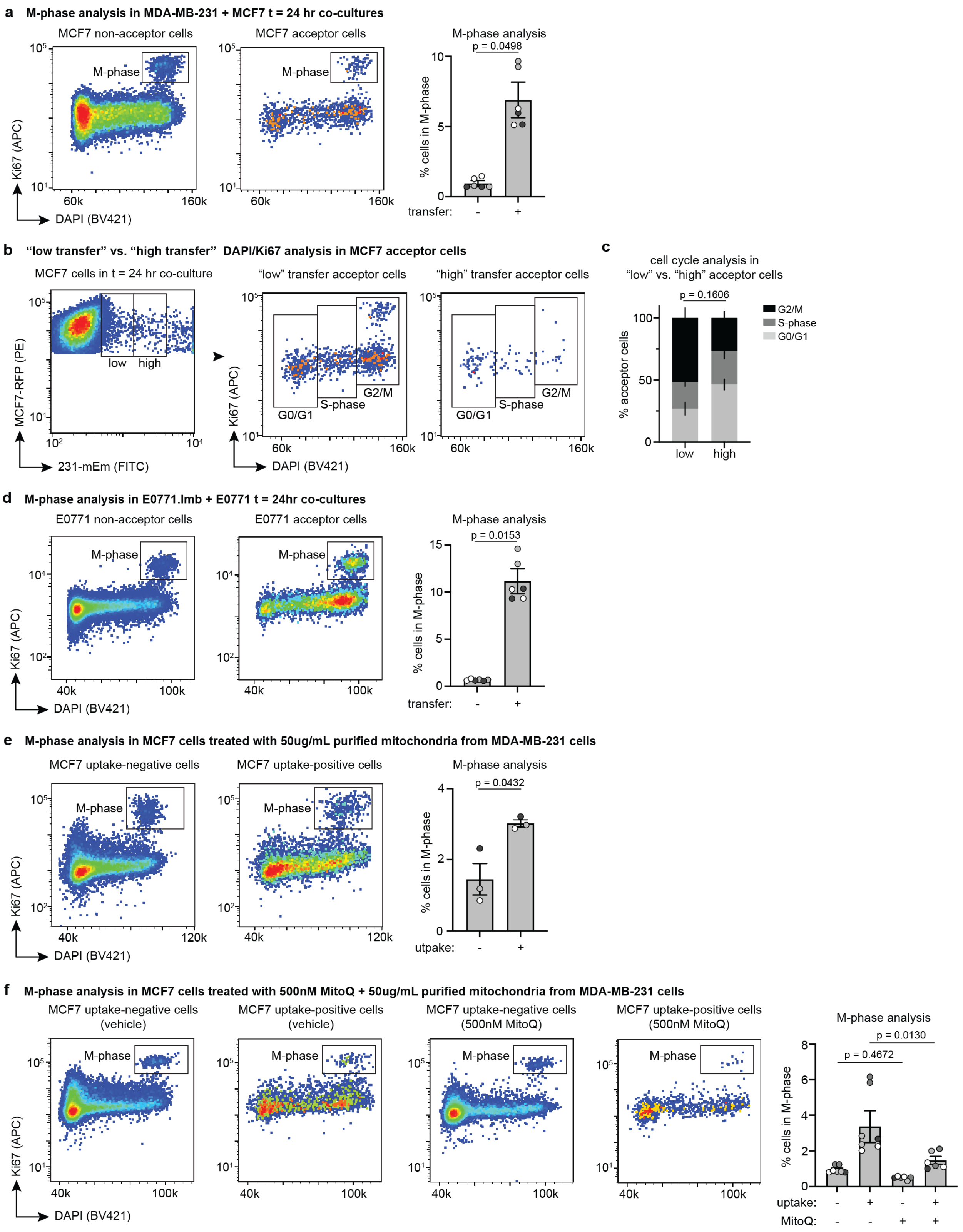


**Supplemental Figure 3. Mitotic phase analysis and “low” vs. “high” mitochondrial transfer acceptors.**

**(a)** Left: Representative flow cytometry plots of non-acceptor and acceptor MCF7 cells in 24 hour co-cultures. A gate is drawn around cells in the M-phase. Right: Quantification of the proportion of non-acceptor and acceptor MCF7 cells in the M-phase. **(b)** Left: Representative gating strategy of “low” and “high” MCF7 acceptor cells. The boundary between “low” and “high” gates was evenly divided between the min. and max. fluorescence intensity of MCF7 acceptor cells. Right: Representative DAPI/Ki67 cell cycle analysis flow cytometry plots in “low” and “high” transfer acceptor cells. Because of the low number of cells in the “high” transfer gate, there is higher variability; however, we provide the cell counts for “low”/”high” populations of each technical replicate: Biological replicate 1, technical replicate A: 2137/90; biological replicate 1, technical replicate B: 2384/98. Biological replicate 2, technical replicate A: 939/94; biological replicate 2, technical replicate B: 929/90. Biological replicate 3, technical replicate A: 1032/109; biological replicate 3, technical replicate B: 877/109. **(c)** Quantification of DAPI/Ki67 cell cycle analysis in “low” and “high” transfer acceptor cells. Statistics are only shown for G2/M-phase comparisons depicted in black. M-phase only analysis is not provided due to the low cell numbers. **(d)** Left: Representative flow cytometry plots of non-acceptor and acceptor E0771 cells in 24 hour co-cultures. A gate is drawn around cells in the M-phase. Right: Quantification of the proportion of non-acceptor and acceptor E0771 cells in the M-phase. **(e)** Left: Representative flow cytometry plots of uptake-negative and uptake-positive MCF7 cells in 24 hour monocultures treated with 50µg/mL of purified mEm+ mitochondria from 231 cells. A gate is drawn around cells in the M-phase. Right: Quantification of the proportion of uptake-negative and uptake-positive MCF7 cells in the M-phase. **(f)** Left: Representative flow cytometry plots of uptake-negative and uptake-positive MCF7 cells in 24 hour monocultures treated with 50µg/mL of purified mEm+ mitochondria from 231 cells and 500nM mitoquinol mesylate (MitoQ) or vehicle. Right: Quantification of the proportion of uptake-negative and uptake-positive MCF7 cells in the M-phase with treatment of 500nM MitoQ or vehicle. For all graphs with data points displayed, each data point represents a technical replicate with each color representing a biological replicate. Paired t-test (**a-e**), two-way ANOVA (Fisher’s LSD post-hoc; **f**).
